## Supplemental Results for "When effort matters: Expectations of reward and efficacy guide cognitive control allocation"

**Supplemental Material**

Table S 1. *Effects of Reward and Efficacy on Performance – Study 1*

| | **Accuracy** | | | **Accurate RT** | | |
|---|---|---|---|---|---|---|
| *Predictors* | *Log-Odds* | *CI* | *p* | *Estimates* | *CI* | *p* |
| (Intercept) | 2.16 | 1.87 – 2.45 | **<0.001** | 600.30 | 581.46 – 619.13 | **<0.001** |
| Efficacy | 0.10 | -0.06 – 0.26 | 0.214 | -14.55 | -20.63 – -8.46 | **<0.001** |
| Reward | 0.05 | -0.11 – 0.21 | 0.543 | -9.81 | -15.89 – -3.73 | **0.002** |
| Congruency n-i | 0.44 | 0.17 – 0.71 | **0.001** | -55.99 | -68.83 – -43.16 | **<0.001** |
| Congruency c-n | 0.54 | 0.29 – 0.78 | **<0.001** | -15.80 | -23.63 – -7.97 | **<0.001** |
| Trial | 0.13 | 0.05 – 0.21 | **0.002** | -1.53 | -4.58 – 1.53 | 0.327 |
| Efficacy: Reward | 0.15 | -0.17 – 0.46 | 0.357 | -9.75 | -21.92 – 2.41 | 0.116 |
| Observations | 6182 | | | 5435 | | |

Note: Statistically significant p-values (< 0.05) are displayed in bold. Congruency (n-i) refers to the comparison between incongruent and neutral Stroop stimulus; Congruency (c-n) refers to the comparison between neutral and congruent Stroop stimulus.

Table S 2. *Effects of Reward and Efficacy on Performance – Study 2*

| | **Accuracy** | | | **Accurate RT** | | |
|---|---|---|---|---|---|---|
| *Predictors* | *Log-Odds* | *CI* | *p* | *Estimates* | *CI* | *p* |
| (Intercept) | 1.86 | 1.65 – 2.08 | **<0.001** | 648.43 | 631.27 – 665.58 | **<0.001** |
| Efficacy | 0.08 | 0.01 – 0.15 | **0.033** | -5.89 | -10.70 – -1.08 | **0.016** |
| Reward | 0.01 | -0.07 – 0.08 | 0.885 | -5.03 | -8.50 – -1.56 | **0.004** |
| Congruency n-i | 0.65 | 0.44 – 0.85 | **<0.001** | -66.17 | -76.64 – -55.69 | **<0.001** |
| Congruency c-n | 0.34 | 0.18 – 0.50 | **<0.001** | -15.23 | -20.63 – -9.83 | **<0.001** |
| Trial | 0.04 | 0.00 – 0.07 | **0.049** | -10.87 | -12.61 – -9.12 | **<0.001** |
| Efficacy: Reward | 0.02 | -0.13 – 0.16 | 0.791 | -9.23 | -16.16 – -2.29 | **0.009** |
| Observations | 24404 | | | 20510 | | |

Note: Statistically significant p-values (< 0.05) are displayed in bold. Congruency (n-i) refers to the comparison between incongruent and neutral Stroop stimulus; Congruency (c-n) refers to the comparison between neutral and congruent Stroop stimulus.

In addition to the within-study effects reported above, we also saw an overall difference in performance between the studies. Relative to Study 1, participants in Study 2 were slower overall ($b = -51.95$, $p <.001$) and less accurate ($b = -0.50$, $p = .013$), which could be attributable to differences in task design or participant pools.

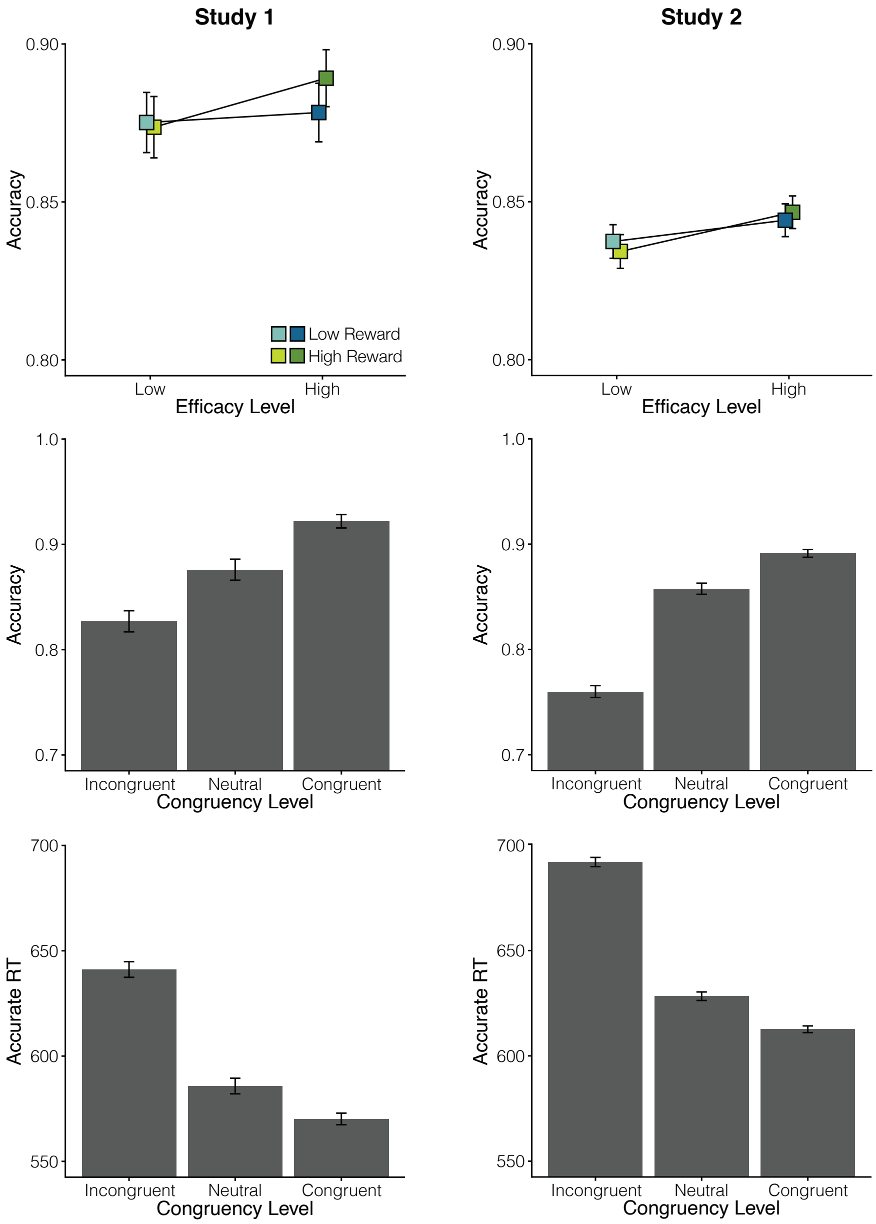


**Figure S 1. Performance effects across both Studies.** Top: Study 1, Bottom: Study 2. **A.** Effects of Reward and Efficacy on performance accuracy. **B.** Effect of Stroop congruency on accuracy. **C.** Effects of Stroop Congruency on accurate RT.

Table S 3. *Between-Study Comparison of Behavioral Effects*

| | **Accuracy** | | | **Accurate RT** | | |
|---|---|---|---|---|---|---|
| *Predictors* | *Log-Odds* | *CI* | *p* | *Estimates* | *CI* | *p* |
| (Intercept) | 2.11 | 1.91 – 2.31 | **<0.001** | 624.25 | 609.99 – 638.51 | **<0.001** |
| Efficacy | 0.09 | 0.00 – 0.17 | **0.047** | -10.26 | -15.01 – -5.51 | **<0.001** |
| Reward | 0.03 | -0.06 – 0.12 | 0.503 | -7.52 | -11.23 – -3.81 | **<0.001** |
| Congruency n-i | 0.53 | 0.35 – 0.72 | **<0.001** | -61.12 | -70.24 – -52.00 | **<0.001** |
| Congruency c-n | 0.45 | 0.29 – 0.60 | **<0.001** | -15.54 | -20.87 – -10.22 | **<0.001** |
| Trial | 0.08 | 0.04 – 0.13 | **<0.001** | -4.01 | -5.99 – -2.03 | **<0.001** |
| Efficacy: Reward | 0.08 | -0.09 – 0.25 | 0.370 | -9.44 | -16.87 – -2.01 | **0.013** |
| Study | -0.50 | -0.89 – -0.11 | **0.013** | 51.95 | 23.43 – 80.47 | **<0.001** |
| Efficacy: Study | -0.02 | -0.19 – 0.15 | 0.816 | 8.61 | -0.88 – 18.10 | 0.075 |
| Reward: Study | -0.05 | -0.23 – 0.12 | 0.560 | 4.97 | -2.46 – 12.40 | 0.190 |
| Congruency n-i : Study | 0.24 | -0.12 – 0.61 | 0.193 | -10.11 | -28.35 – 8.13 | 0.277 |
| Congruency c-n : Study | -0.23 | -0.53 – 0.07 | 0.140 | 0.66 | -10.00 – 11.31 | 0.904 |
| Trial : Study | -0.13 | -0.22 – -0.03 | **0.009** | -4.53 | -8.48 – -0.57 | **0.025** |
| Efficacy: Reward:Study | -0.12 | -0.47 – 0.22 | 0.482 | 0.42 | -14.45 – 15.28 | 0.956 |
| N | 65 SubID | | | 65 SubID | | |
| Observations | 30566 | | | 25945 | | |

Note: Statistically significant p-values (< 0.05) are displayed in bold. Congruency (n-i) refers to the comparison between incongruent and neutral Stroop stimulus; Congruency (c-n) refers to the comparison between neutral and congruent Stroop stimulus.

Table S 4. *Parametric Effects of Reward and Efficacy on Performance – Study 3*

| | **Accuracy** | | | **Accurate RT** | | |
|---|---|---|---|---|---|---|
| *Predictors* | *Log-Odds* | *CI* | *p* | *Estimates* | *CI* | *p* |
| (Intercept) | 1.78 | 1.51 – 2.04 | **<0.001** | 602.99 | 588.27 – 617.71 | **<0.001** |
| Efficacy | -0.01 | -0.06 – 0.04 | 0.746 | -3.85 | -6.10 – -1.60 | **0.001** |
| Reward | 0.03 | -0.01 – 0.08 | 0.167 | -7.02 | -9.24 – -4.81 | **<0.001** |
| Congruency n-i | 0.28 | 0.13 – 0.42 | **<0.001** | -52.34 | -66.37 – -38.31 | **<0.001** |
| Congruency c-n | 0.55 | 0.37 – 0.73 | **<0.001** | -18.15 | -24.81 – -11.48 | **<0.001** |
| Trial | 0.08 | 0.03 – 0.14 | **0.003** | -4.39 | -6.88 – -1.90 | **0.001** |
| Efficacy:Reward | 0.01 | -0.03 – 0.05 | 0.621 | -2.27 | -4.27 – -0.26 | **0.027** |
| N | 35 | | | 35 | | |
| Observations | 10251 | | | 8531 | | |

Note: Statistically significant p-values (< 0.05) are displayed in bold. Congruency (n-i) refers to the comparison between incongruent and neutral Stroop stimulus; Congruency (c-n) refers to the comparison between neutral and congruent Stroop stimulus.

Table S 5. *ERPs predict behavior within incentive conditions*

| | Accuracy | | | Accurate RT | | |
|---|---|---|---|---|---|---|
| *Predictors* | *Log-Odds* | *CI* | *p* | *Estimates* | *CI* | *p* |
| (Intercept) | 1.87 | 1.65 – 2.08 | **<0.001** | 647.64 | 630.60 – 664.68 | **<0.001** |
| Efficacy | 0.08 | 0.01 – 0.16 | **0.034** | -3.79 | -8.62 – 1.04 | 0.124 |
| Reward | 0.03 | -0.05 – 0.10 | 0.455 | -2.99 | -6.59 – 0.61 | 0.104 |
| Congruency n-i | 0.65 | 0.44 – 0.85 | **<0.001** | -63.99 | -68.85 – -59.13 | **<0.001** |
| Congruency c-n | 0.34 | 0.18 – 0.49 | **<0.001** | -15.82 | -20.23 – -11.41 | **<0.001** |
| Baseline | 0.02 | -0.02 – 0.07 | 0.333 | 2.24 | 0.05 – 4.43 | **0.045** |
| Trial | 0.04 | 0.00 – 0.08 | **0.039** | -10.79 | -12.60 – -8.98 | **<0.001** |
| Efficacy: Reward | 0.01 | -0.14 – 0.16 | 0.905 | -9.85 | -17.04 – -2.66 | **0.007** |
| Efficacy$_l$ : Reward$_l$ : P3b | 0.05 | -0.03 – 0.12 | 0.223 | -11.33 | -15.05 – -7.62 | **<0.001** |
| Efficacy$_h$ : Reward$_l$ : P3b | 0.16 | 0.08 – 0.24 | **<0.001** | -4.89 | -8.50 – -1.28 | **0.008** |
| Efficacy$_l$ : Reward$_h$ : P3b | -0.01 | -0.09 – 0.07 | 0.801 | -4.53 | -8.23 – -0.84 | **0.016** |
| Efficacy$_h$ : Reward$_h$ : P3b | 0.10 | 0.03 – 0.18 | **0.009** | -7.52 | -11.11 – -3.94 | **<0.001** |
| Efficacy$_l$ : Reward$_l$ : CNV | -0.06 | -0.14 – 0.02 | 0.117 | 16.50 | 11.72 – 21.28 | **<0.001** |
| Efficacy$_h$ : Reward$_l$ : CNV | -0.08 | -0.16 – -0.00 | **0.045** | 13.28 | 8.50 – 18.05 | **<0.001** |
| Efficacy$_l$ : Reward$_h$ : CNV | -0.10 | -0.18 – -0.03 | **0.007** | 13.89 | 9.10 – 18.68 | **<0.001** |
| Efficacy$_h$ : Reward$_h$ : CNV | -0.15 | -0.23 – -0.07 | **<0.001** | 18.68 | 13.96 – 23.41 | **<0.001** |
| N | 44 | | | 44 | | |
| Observations | 22580 | | | 18999 | | |

### Pupil dilation tracks uncertainty-linked arousal, not proactive control

Table S 6. *Incentive effects on pupil dilation*

| | Pupil Response | | |
|---|---|---|---|
| *Predictors* | *Estimates* | *CI* | *p* |
| (Intercept) | 0.28 | 0.22 – 0.35 | **<0.001** |
| Efficacy | -0.06 | -0.08 – -0.04 | **<0.001** |
| Reward | 0.01 | -0.01 – 0.03 | 0.243 |
| Trial | -0.19 | -0.28 – -0.10 | **<0.001** |
| Efficacy * Reward | -0.02 | -0.05 – 0.02 | 0.309 |
| Observations | 19986 | | |

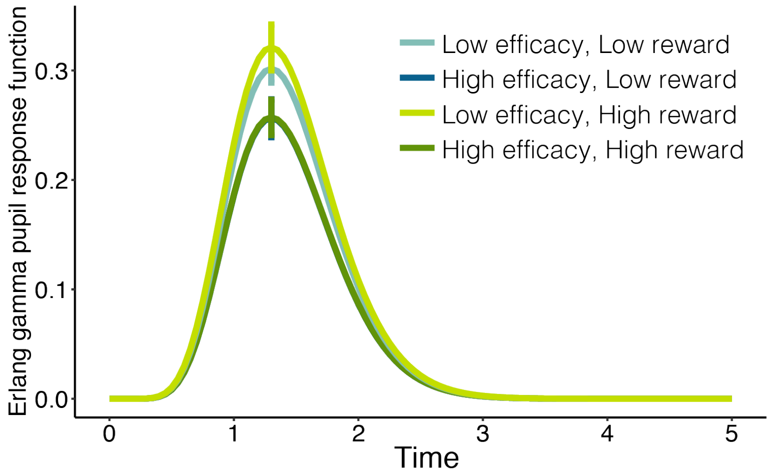


**Figure S 2. Pupillary response increases with uncertainty, not motivation.** Cue-related pupillary response estimated using deconvolution as a function of incentive condition. Error bars represent the 95% CI of the scaling parameters.

**Response-related signals reflect violations of performance criteria and expectations**

We tested how incentives would modulate response and outcome evaluation. To index response evaluation, we analyzed the error related negativity (ERN). This signal is sensitive to response errors and has been proposed to reflect reward prediction errors computed based on the change in expected outcome as a function of internal evaluations of the response (Holroyd & Coles, 2002). In our task, performance was rewarded if participants were both fast and accurate. Hence, internal response evaluations should be sensitive to both the accuracy and speed of the submitted response, with larger ERN amplitudes for both incorrect and slower responses. Consistent with this assumption, we found additive effects of response accuracy and RT (Table S5). ERN amplitude was larger for errors compared to correct responses, and increased with increasing response time. In addition to the linear increase of ERN amplitude with RT, we also observed a quadratic component, such that RT effects tapered off around the time of the criterion (~750 ms, cf. Fig. S3). In addition, we tested whether participants were sensitive to incentive-dependent expectations about their response accuracy. To do so, we computed the average accuracy within each incentive condition (probability correct for each combination of efficacy and reward). We found that higher incentive-dependent accuracy expectations, amplified accuracy effects on the ERN (Fig. S3). Congruency effects on ERN amplitude were modulated by RT, such that differences in ERN as a function of congruency conditions washed out for longer RTs (Fig. S3). As reported in the main manuscript, when testing our prediction that higher reward and efficacy would amplify internal monitoring, we found the expected 3-way interaction between efficacy, reward, and accuracy (Table S5). However, at odds with our prediction, the effect was driven by largest ERN amplitude on error trials in the condition in which both efficacy and reward were low (Table S6), and this pattern was suspiciously mirrored in error RTs (Table S 7 and Fig. S3). While this pattern of activity was not predicted (we predicted largest ERN amplitudes to errors on high reward, high efficacy trials), it is notable that it mirrors the pattern we observed for error RTs (Fig. S3), where low-reward/low-efficacy trials were associated with the fastest errors (consistent with effort minimization by, e.g., responding with any key; $b = 29.08$, $p = .001$; Table S7). Speculatively, these findings may reflect a shift from more effortful proactive to less effortful reactive control as the expected value of proactive control decreased (Braver, 2012). Thus, participants may have responded more impulsively, and then evaluated those responses immediately after they were executed. On the large proportion of incongruent error trials participants may further have re-evaluated the value of control in a reactive manner given the new information about the difficulty of the trial. Future research will need to test specific predictions of this ad-hoc interpretation in a principled manner. Given the sensitivity of the ERN to multiple aspects of performance and performance expectations, it is possible that the complex interaction pattern reflects different mixtures of these multiple factors across incentive conditions.

Table S 7. *Reward, Efficacy, and Performance Effects on ERN*

| | **ERN** | | |
|---|---|---|---|
| *Predictors* | *Estimates* | *CI* | *p* |
| (Intercept) | 2.95 | 1.19 – 4.71 | **0.001** |
| Efficacy | 0.09 | -0.14 – 0.33 | 0.444 |
| Reward | 0.03 | -0.20 – 0.26 | 0.773 |
| Accuracy | 4.31 | 3.56 – 5.07 | **<0.001** |
| Congruency n-i | 7.87 | 4.00 – 11.73 | **<0.001** |
| Congruency c-n | -4.76 | -8.35 – -1.17 | **0.009** |
| RT | -7.59 | -12.48 – -2.71 | **0.002** |
| RT quadratic | 4.57 | 0.81 – 8.34 | **0.017** |
| Mean Accuracy (mAcc) | -1.96 | -5.02 – 1.11 | 0.210 |
| Baseline | -0.36 | -0.38 – -0.35 | **<0.001** |
| Efficacy: Reward | -0.62 | -1.08 – -0.15 | **0.009** |
| Efficacy: Accuracy | -0.07 | -0.54 – 0.40 | 0.755 |
| Reward: Accuracy | -0.36 | -0.82 – 0.10 | 0.127 |
| Congruency n-i : RT | -19.62 | -31.37 – -7.88 | **0.001** |
| Congruency c-n : RT | 11.54 | 0.38 – 22.70 | **0.043** |
| Congruency n-i : RT qu. | 11.50 | 2.80 – 20.20 | **0.010** |
| Congruency c-n : RT qu. | -6.28 | -14.73 – 2.16 | 0.145 |
| Accuracy : mAcc | 7.67 | 2.24 – 13.11 | **0.006** |
| Efficacy: Reward: Accuracy | 1.52 | 0.60 – 2.45 | **0.001** |
| $N_{SubID}$ | 44 | | |
| Observations | 22954 | | |

Note: Statistically significant p-values (< 0.05) are displayed in bold. Congruency (i-n) refers to the comparison between incongruent and neutral Stroop stimulus; Congruency (n-c) refers to the comparison between neutral and congruent Stroop stimulus.

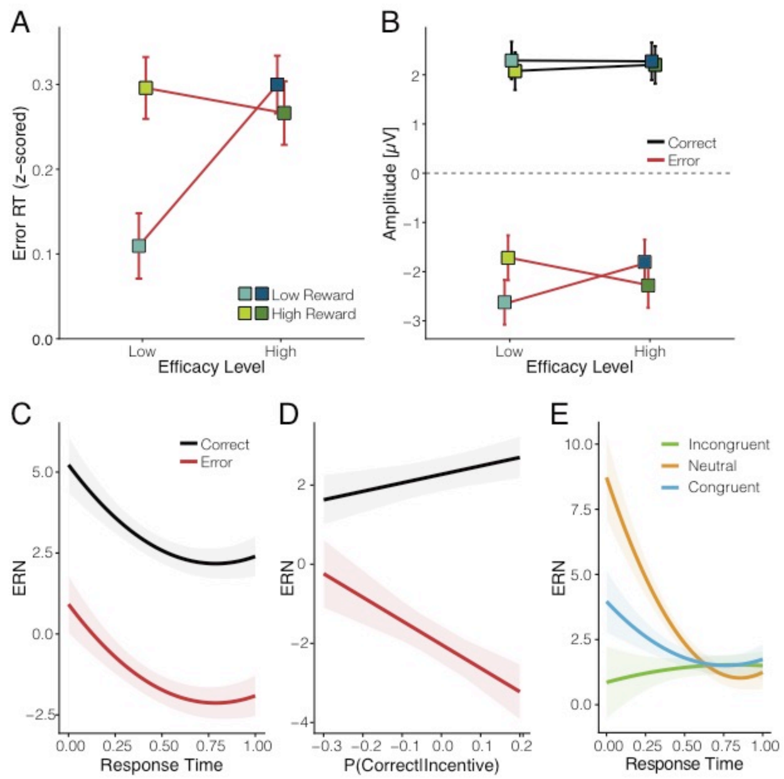


**Figure S 3. Response-evaluation is reflects violations of performance expectations, not incentive-driven outcome salience. A.** Error RTs are plotted as a function of incentive conditions and show the same pattern as ERN amplitudes. **B.** Response-locked amplitudes are plotted as a function of response accuracy and incentive conditions. ERN amplitude is largest on error trials with low reward and low efficacy. **C.** Fixed effects of accuracy and response time on response- locked activity. Response-locked amplitudes are more negative for errors, as well as slower responses. **D.** Fixed effects of accuracy in interaction with incentive-wise mean accuracy. Amplitudes for correct and incorrect responses are more similar when errors are more frequent. Error-related activity is more negative when errors are less frequent. **E**. Fixed effects of Congruency in interaction with RT. Congruency effects on the ERN wash out for longer response times. Shaded and regular error bars represent standard error of the mean.

Table S 8. *Incentive and Performance Effects on ERN Amplitude in Correct and Error Trials*

| | **ERN error** | | | **ERN correct** | | |
|---|---|---|---|---|---|---|
| *Predictors* | *Estimates* | *CI* | *p* | *Estimates* | *CI* | *p* |
| (Intercept) | 2.39 | -1.66 – 6.43 | 0.248 | 4.48 | 2.54 – 6.42 | **<0.001** |
| Efficacy | 0.12 | -0.32 – 0.57 | 0.582 | 0.05 | -0.13 – 0.24 | 0.581 |
| Reward | 0.19 | -0.24 – 0.63 | 0.382 | -0.14 | -0.33 – 0.04 | 0.120 |
| Congruency2-1 | -2.38 | -11.63 – 6.88 | 0.615 | 10.56 | 6.29 – 14.83 | **<0.001** |
| Congruency3-2 | -2.09 | -11.21 – 7.03 | 0.653 | -5.50 | -9.42 – -1.59 | **0.006** |
| RT | -13.53 | -25.21 – -1.85 | **0.023** | -5.54 | -10.92 – -0.16 | **0.044** |
| RT quadratic | 9.66 | 1.08 – 18.24 | **0.027** | 2.93 | -1.22 – 7.08 | 0.167 |
| meanCorr1 | -5.42 | -10.74 – -0.09 | **0.046** | 1.57 | -1.46 – 4.61 | 0.310 |
| Baseline | -0.37 | -0.40 – -0.34 | **<0.001** | -0.36 | -0.37 – -0.35 | **<0.001** |
| Efficacy: Reward | -1.34 | -2.21 – -0.47 | **0.003** | 0.15 | -0.22 – 0.51 | 0.432 |
| Congruency2-1 * RT | 8.12 | -19.67 – 35.92 | 0.567 | -27.30 | -40.31 – -14.29 | **<0.001** |
| Congruency3-2 * RT | 3.95 | -23.93 – 31.82 | 0.781 | 13.81 | 1.60 – 26.02 | **0.027** |
| Congruency2-1 * RT qu. | -7.08 | -27.44 – 13.27 | 0.495 | 16.87 | 7.21 – 26.54 | **0.001** |
| Congruency3-2 * RT qu. | -0.41 | -21.16 – 20.34 | 0.969 | -8.07 | -17.34 – 1.20 | 0.088 |
| N | 44 SubID | | | 44 SubID | | |
| Observations | 3640 | | | 19314 | | |

Note: Statistically significant p-values (< 0.05) are displayed in bold. Congruency (i-n) refers to the comparison between incongruent and neutral Stroop stimulus; Congruency (n-c) refers to the comparison between neutral and congruent Stroop stimulus.

Table S 9. *Incentive Effects on Error RT*

| | **Error RT** | | |
|---|---|---|---|
| *Predictors* | *Estimates* | *CI* | *p* |
| (Intercept) | 682.12 | 660.66 – 703.57 | **<0.001** |
| Reward | 9.45 | 1.20 – 17.70 | **0.025** |
| Congruency n-i | -47.70 | -60.83 – -34.57 | **<0.001** |
| Congruency c-n | 10.16 | -8.41 – 28.74 | 0.284 |
| Trial | -10.11 | -14.22 – -6.00 | **<0.001** |
| Reward [l] : Efficacy | 29.08 | 14.24 – 43.92 | **<0.001** |
| Reward[h] : Efficacy | -4.51 | -19.33 – 10.31 | 0.551 |
| N SubID | 44 | | |
| Observations | 3875 | | |

Note: Statistically significant p-values (< 0.05) are displayed in bold. Congruency (i-n) refers to the comparison between incongruent and neutral Stroop stimulus; Congruency (n-c) refers to the comparison between neutral and congruent Stroop stimulus.

In addition to ERN amplitude, we analyzed response-locked theta power, which indexes similar, although not functionally identical processes to the ERN (Beatty, Buzzell, Roberts, & McDonald, 2020). We had predicted that midfrontal theta following the response would be largest when participants make errors on high reward and efficacy trials. Our results (Table S8) are partially in line with our predictions, such that theta power was higher when participants made errors on high efficacy trials, reflected in a significant interaction of efficacy and accuracy (b = -276.12, p = .013), as well reliably higher theta power for high compared to low efficacy on error, but not correct trials (error: b = 318.45, p = .002, correct: b = 42.33, p = .337). However, we observed no reliable effects of reward or interactions with reward and if anything theta power was lower on high relative to low reward trials.

Table S 10. *Incentive Effects on Error Processing in Midfrontal Theta*

| | **Theta Power** | | |
|---|---|---|---|
| *Predictors* | *Estimates* | *CI* | *p* |
| (Intercept) | 4337.09 | 3912.50 – 4761.68 | **<0.001** |
| Efficacy | 180.39 | 71.12 – 289.66 | **0.001** |
| Reward | -54.11 | -163.00 – 54.77 | 0.330 |
| Accuracy | -1529.32 | -1935.98 – -1122.66 | **<0.001** |
| Baseline | 212.27 | 169.27 – 255.28 | **<0.001** |
| Efficacy: Accuracy | -276.12 | -494.68 – -57.57 | **0.013** |
| Reward: Accuracy | 166.43 | -51.34 – 384.20 | 0.134 |
| Observations | 23251 | | |

Note: Statistically significant p-values (< 0.05) are displayed in bold

**Feedback evaluation reflects outcome magnitude and predictability**

We had predicted that incentives would shape feedback processing, such that reward effects would be amplified under high efficacy. To test this prediction, we quantified the magnitude of the FRN peak-to-peak relative to the preceding positive deflection. This measure eliminates baseline confounds that may lead to spurious effects (e.g. due to anticipatory signals). Consistent with the reward prediction error account of the FRN (Holroyd & Coles, 2002), the peak-to-peak FRN reflected reward receipt versus omission, with larger FRN amplitudes for unrewarded compared to rewarded trials (b = 0.80, p <.001), as well as stronger effects for larger rewards (b = 0.81, p = .007). However, in addition, this measure revealed an interaction of reward receipt with Efficacy, such that reward effects were reduced for high compared to low efficacy (b = -0.83, p = .007). While at odds with our original prediction, this finding is plausible, given that under high efficacy rewards were controllable, and could be predicted based on performance (cf. response evaluation), thus, the impact of these rewards was reduced.

Table S 11. *Incentive Modulations of Reward Processing (FRN)*

| | **Peak to peak FRN amplitude** | | |
|---|---|---|---|
| *Predictors* | *Estimates* | *CI* | *p* |
| (Intercept) | -34.97 | -37.29 – -32.64 | **<0.001** |
| Is Rewarded y-n | 0.80 | 0.41 – 1.19 | **<0.001** |
| Efficacy | -0.00 | -0.30 – 0.30 | 0.996 |
| Reward | 0.07 | -0.23 – 0.37 | 0.655 |
| Congruency n-i | -0.08 | -0.46 – 0.30 | 0.691 |
| Congruencyc-n | 0.09 | -0.27 – 0.44 | 0.631 |
| Trial | -0.52 | -0.66 – -0.38 | **<0.001** |
| Is Rewarded y-n:Efficacy | -0.83 | -1.43 – -0.23 | **0.007** |
| Is Rewarded y-n:Reward | 0.81 | 0.22 – 1.41 | **0.007** |
| Observations | 22721 | | |

Note: Statistically significant p-values (< 0.05) are displayed in bold. Congruency (n-i) refers to the comparison between incongruent and neutral Stroop stimulus; Congruency (c-n) refers to the comparison between neutral and congruent Stroop stimulus.

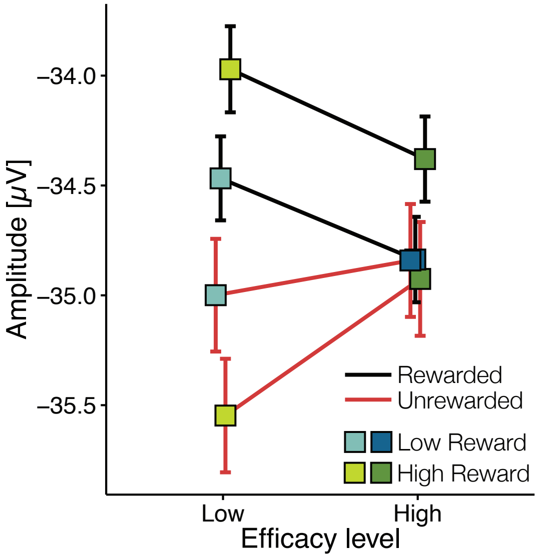


**Figure S 4. Outcome effects on FRN amplitude are larger for larger rewards, but lower efficacy.** Mean peak-to-peak FRN amplitude as a function of reward receipt vs. omission, reward magnitude, and efficacy on FRN amplitude. Error bars represent standard errors of the mean.

**Supplementary Discussion**

Our ERN results - largest amplitudes on error trials with low reward and low efficacy - are inconsistent with a motivational salience account. Instead we found response-locked activity that tracked violations of either performance criterion (accuracy or response speed): an ERN to errors compared to correct responses, as well as more negative amplitudes to longer RTs, maximal around the time of the reward deadline (Fig. S3, Table S4). ERN amplitudes were also larger when errors were less expected based on one's average performance in a given incentive condition (i.e., when performing well overall) (Brown & Braver, 2005; Hughes & Yeung, 2011). Our results at the time of feedback (FRN) - reduced effects of reward receipt vs omission under high efficacy - are also inconsistent with a motivational salience account. Instead, they indicate that predictability modulated outcome evaluation. Under high (but not low) efficacy, outcome information is redundant to the degree that participants can internally evaluate their performance, as suggested by our ERN results and previous work (Bellebaum & Colosio, 2014; Bultena, Danielmeier, Bekkering, & Lemhöfer, 2017; Frömer, Nassar, Stürmer, Sommer, & Yeung, 2018; Holroyd & Coles, 2002), reducing the response to feedback. As difficulty manipulations are known to influence such internal evaluations (Boldt, de Gardelle, & Yeung, 2017), our results call for caution when interpreting corresponding increases (Hernandez Lallement et al., 2014; Ma, Meng, Wang, & Shen, 2014; Wang, Zheng, & Meng, 2017), or decreases (effort discounting; Botvinick, Huffstetler, & McGuire, 2009) in outcome valuation as effort-related. Taken together, the observed sensitivity of ERN and FRN to surprising evaluative information is in line with the reward prediction error account of the ERN and FRN(Holroyd & Coles, 2002), and emphasizes the interaction of internal performance monitoring and feedback processing(Frömer et al., 2018). Thus, in contrast to the robust incentive-driven increase in markers of proactive control, our results are inconsistent with incentive-driven increases in motivational salience of outcomes, reactive control, or overall engagement.

With that said, there was little use for incentive-based reactive control in the present paradigm, as feedback was only informative about the success on the current, but not the value of effort on the subsequent trial. This difference in functional utility of feedback may explain why our results square with previously reported larger FRN amplitudes for more reliable feedback. When participants were led to believe that some equally unreliable feedback could be used to improve task performance – i.e. that reactive control would be more efficacious – their FRN amplitudes to that feedback were larger (Muhlberger, Angus, Jonas, Harmon-Jones, & Harmon-Jones, 2017; Schiffer, Siletti, Waszak, & Yeung, 2017). Thus, effort cost-benefit analyses, determining the intensity *and* type of the optimal control signal, may yield different results for proactive and reactive control signals, depending on the task (Braver, 2012; Shenhav, Botvinick, & Cohen, 2013; Shenhav, Cohen, & Botvinick, 2016). Thus, additional work is needed to investigate under which conditions and how reward and efficacy shape reactive trial-to-trial adjustments of control.

**Supplementary Method**

Table S 12. *Performance and Reward Summary Statistics*

| | Fast and accurate | Rewarded |
|---|---|---|
| **Study 1** | M = 0.77, SD = 0.10 | M = 0.76, SD = 0.10 |
| **Study 2** | M = 0.61, SD = 0.16 | M = 0.60; SD = 0.15 |

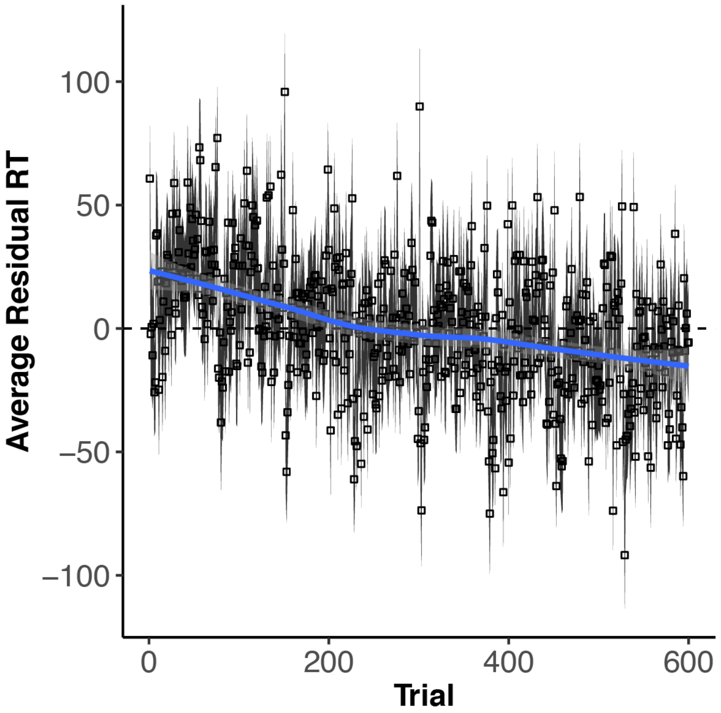


**Figure S 5. Residuals from model without trial regressors reveal approximately linear changes over the course of the experiment.** Plotted are the average residuals for each trial in Study 2 with standard errors as shaded error bars. Superimposed is an unconstrained spline fit (blue line) that yields an approximately linear trend.
